# Microglia and its P2RY12 Receptors Regulate Seizure Severity

**DOI:** 10.64898/2025.12.11.693689

**Authors:** Leigh Ellen Fremuth, Synphane Gibbs-Shelton, Madison Doceti, Aida Oryza Lopez-Ortiz, Abigayle Duffy, Ronald P. Gaykema, Kiran Singh, Audrey Nguyen, Sophia Cheng, Edward Perez-Reyes, Ukpong B. Eyo

## Abstract

Microglia have emerged as possible regulators of seizures with previous approaches employed being insufficiently selective of microglial-specific contributions. To more definitely determine microglial roles in seizure severity, we used the recently developed genetic microglial-deficient *Csf1r^ΔFIRE/ΔFIRE^* (FIRE KO) mouse model. In this model, mice lack microglia but retain brain border associated macrophages (BAMs). Using two experimental seizure paradigms with either a chemoconvulsant or electrical stimulation, we confirm that a microglial deficiency exacerbates seizures, reduces survival and facilitates the likelihood of developing spontaneous recurrent seizures, indicating that microglia constrain seizure activity across different seizure paradigms. To gain insights into molecular regulators of seizure severity under microglial regulation, we examined P2RY12 contributions and demonstrate that a loss of microglial P2RY12 increased seizure severity in both global and microglial-specific knockout mice indicating that microglia suppress seizure severity in chemoconvulsive models through P2RY12 signaling. During seizures, P2RY12-deficient microglia displayed altered process complexity, accompanied by increased neuronal activation and reduced inhibitory tone. These results link impaired microglial responses to heightened seizure susceptibility and network excitability. Together, our findings establish microglia as protective regulators of seizure activity and identify P2RY12 signaling as a critical mediator of microglia protective actions.

## INTRODUCTION

Statistics indicate that about 1 in 26 people will develop epilepsy in their lifetime^1–3^ making it a common disorder. Epilepsy is characterized by the occurrence of seizures as its most obvious manifestation. While seizures are known to result from an imbalance in neuronal excitation and inhibition, treatment approaches that target neuron-autonomous mechanisms have remained insufficient in treating seizure disorders in many patients^4–6^ raising the need for identifying alternative treatment approaches/targets that are relevant for seizure induction, maintenance and/or termination. Of particular interest, glial cells that modulate neuronal activity are receiving increasing attention as important contributors to seizures and epilepsy^7–9^.

Emerging findings indicate that inflammatory processes contribute to seizures^10–12^. However, cell-specific assessments of neuroinflammatory contributions to seizures require further research. Among brain cells, microglia are uniquely positioned at the interface of the nervous and immune systems being the primary immunocompetent cells of the brain. Our understanding of microglial roles in seizures and epilepsy development is still at its infancy. Results indicate complex roles for microglia with some studies suggesting seizure-limiting^13–17^ and others suggesting seizure-promoting roles^18–24^. However, appropriate tools are required for precise investigation and the elucidation of relevant, targetable mechanisms.

Recently, a powerful tool for microglial studies was developed in the form of the generation of Csf1r^ΔFIRE/ΔFIRE^ (FIRE knockout or KO) mouse, where deletion of the fms-intronic regulatory element (FIRE) at the *Csf1r* locus results in the prevention of microglial development without affecting other brain macrophage populations^25^. This mouse model, therefore, provides a powerful microglial-selective model to assess microglial roles in experimental seizure contexts. Interestingly, a recent study suggested that synaptic maturation and seizure susceptibility were unaltered in FIRE KO mice during development^26^. This finding is not congruent with previous literature that suggested microglia play active and protective roles during status epilepticus and during the chronic phase of spontaneous recurring seizures in mice^15^ and a zebrafish developmental Dravet syndrome model of genetic seizures^17^. Moreover, the FIRE KO mice have not been used to interrogate microglial contribution to experimental seizures in adulthood.

Furthermore, whereas some studies define a protective role of the microglia in restricting the severity of seizures through various seizure models, identifying molecular mediators controlling microglial seizure-regulating activities is important for developing therapeutic interventions. Amongst candidates, the P2RY12 receptor is a microglial signature gene^27^ and regulates microglial-neuronal communication through sensing extracellular ADP during neuronal activity from hyperactive neurons to facilitate restoration to homeostatic states ^28–34^. Lack of P2RY12 has been implicated in defective microglial chemotaxis^35^, a defective synaptic management^36,37^, and increased network excitability^38,39^, implying that the P2RY12 may provide a link of mechanisms between microglial surveillance, microglial-neuronal interactions and seizure inhibition.

In this study, we interrogate microglial and P2RY12-specific roles in experimental seizures. First, with FIRE KO mice, we establish a neuroprotective role for microglia in modulating seizure severity in both chemoconvulsive (kainic acid–induced) seizures and electrical kindling seizure paradigms. Second, using P2RY12 global and constitutive microglial knockout lines, we identify a role for microglial P2RY12 in the seizure-related suppression attributed to microglia. Finally, we correlate this role with alterations in microglia process complexity, neuronal cFos activity, and inhibitory VGAT synaptic expression. Thus, we highlight alterations by which microglial P2RY12 preserve network integrity with seizures.

## RESULTS

### “FIRE” KO mice show exacerbated chemoconvulsant seizures

In previous work, we used a pharmacological microglial depletion model to assess microglial contributions to seizures in three seizure paradigms including chemoconvulsive, electrical, and temperature induced febrile seizure models. In those paradigms, we reported exacerbated seizures in microglia with pharmacological microglial elimination^15^. However, pharmacological inhibition of CSF1R using PLX3397, while efficiently eliminating microglia, also targets brain border associated macrophages (BAMs) leaving the possibility that effects seen with that pharmacological approach may be a result of these BAMs. More selective approaches are therefore needed to ascertain microglial roles in experimental seizure paradigms.

To investigate the specific role of microglia in seizure regulation, we utilized Csf1r^ΔFIRE/ΔFIRE^ (hereafter referred to as “FIRE KO”) mice, a genetically engineered model that lacks brain microglia due to deletion of the fms-intronic regulatory element (FIRE) within the *Csf1r* gene^25^. Unlike pharmacological depletion approaches^40^ which affect BAM populations^40,41^, the FIRE KO model provides a selective and permanent loss of microglia while preserving brain border-associated macrophages (BAMs) densities^25^. Given that FIRE KO mice lack microglia but retain other brain macrophage populations, this model allows us to isolate the role of microglia specifically, independent of brain BAMs, in modulating seizure severity. Using this approach, we examined seizure susceptibility across two seizure paradigms in the adult brain.

We crossed FIRE KO mice with CX3CR1-GFP mice to generate CX3CR1^GFP/+^ littermates that would allow for the visualization of myeloid cells by GFP. We confirmed the absence of microglia in the brains of FIRE mice compared to wildtype (FIRE WT) and FIRE heterozygous (FIRE HET) littermate mice (**Fig. 1a-c**). The quantification of GFP^+^ microglia between FIRE WT and FIRE HET was indistinguishable, while FIRE KO had no microglia (**Fig. 1d**), consistent with previous reports^25^. The only GFP^+^ cell detected in FIRE KO mice were border associated macrophages. The selective microglia deficiency in FIRE KO mice allowed us to investigate the impact of microglial absence on chemoconvulsive seizure severity.

**Fig. 1:**
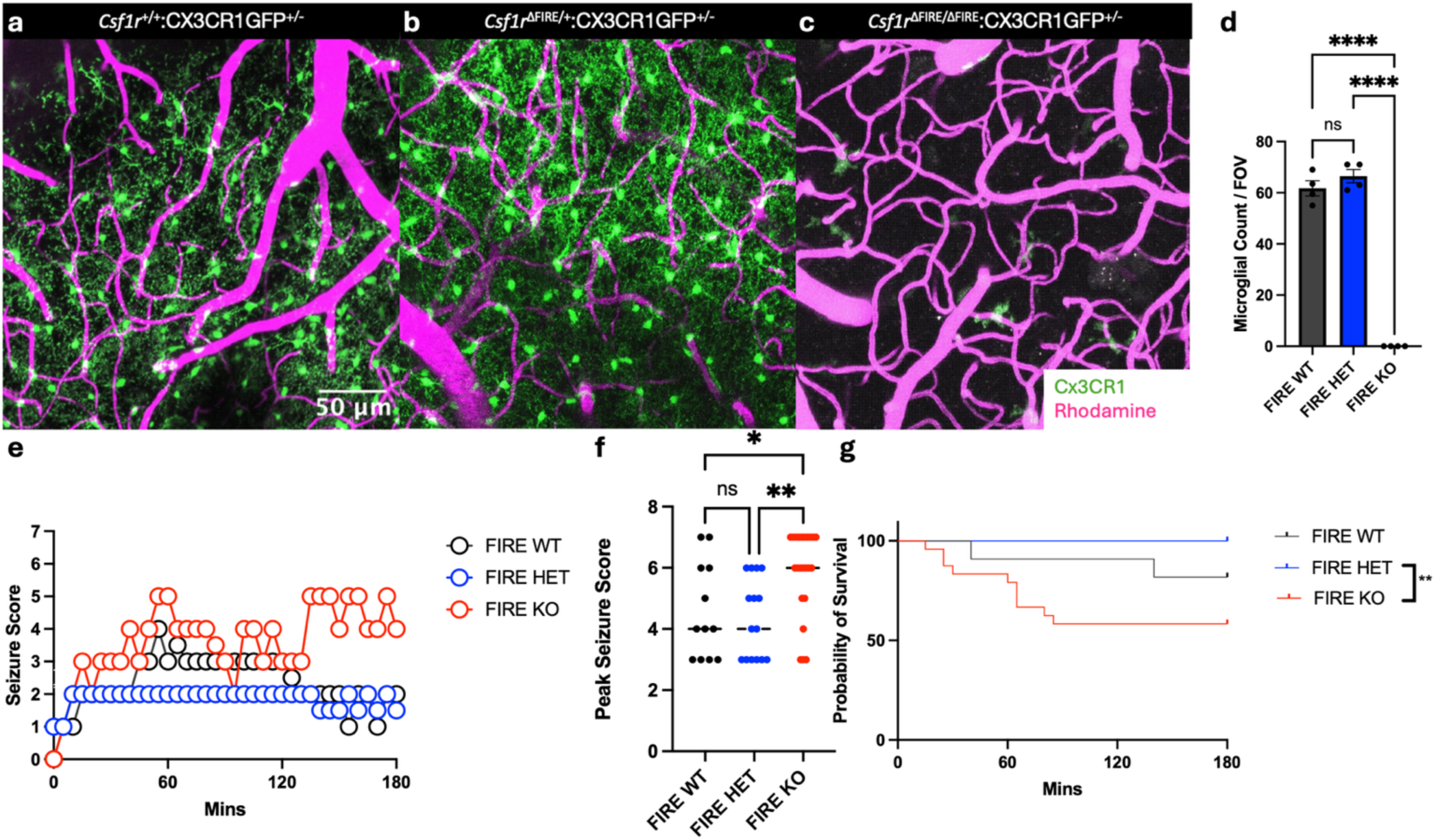
FIRE knockout mice are more susceptible to kainic acid-induced seizures. **a-c**, Representative *in vivo* two photon images showing microglial depletion in Csf1r^ΔFIRE/ΔFIRE^ (FIRE KO) mice compared to wildtype (FIRE WT) and Csf1r^ΔFIRE/+^ (FIRE HET) mice. CX3CR1^GFP/+^ (green) marks microglia, while rhodamine (magenta) highlights the vasculature. **d**, Quantification of microglial numbers in littermate Csf1r^+/+^ (FIRE WT), Csf1r^ΔFIRE/+^ (FIRE HET), and Csf1r^ΔFIRE/ΔFIRE^ (FIRE KO) mice. **e-g**, Seizure severity scores along a modified Racine scale over time (**e**) and peak seizure scores (**f**) in FIRE WT, FIRE HET, and FIRE KO littermate mice following KA induced seizures. **g**, Kaplan-Meier survival curves of FIRE WT, FIRE HET, and FIRE KO littermates that underwent KA-induced seizures. N = 12-23 per group. Error bars represent SEM in **d**. Statistical significance was determined using ANOVA and unpaired t-test. ** p < 0.01; **** p < 0.0001.

Chemoconvulsive seizures were induced by kainic acid (KA) treatment (IP, 24 mg/kg), and seizure severity was quantified using a modified Racine seizure scoring^22^ scheme over time. FIRE KO mice exhibited persistently high seizure scores during the 3h monitoring period and significantly increased peak seizure scores when compared to WT and HET controls (**Fig. 1e-f**). This suggests a heightened susceptibility to chemoconvulsive seizures in the absence of microglia. Moreover, animal survival analysis revealed a reduced probability of survival in FIRE KO mice following seizure induction when compared to HET littermates (**Fig. 1g**), highlighting a critical role for microglia in mitigating seizure-induced mortality. These findings support the hypothesis that microglia play an essential neuroprotective role during seizures by maintaining neural homeostasis, as we and others have previously suggested ^14,15,17,34,42^.

Because previous studies^15^ showed similar results using a pharmacological approach to eliminate microglia with PLX3397^43^ that targets both microglia and brain BAMs^44^, it is possible that these macrophages that persist in FIRE KO mice also contribute to seizure susceptibility to KA. To evaluate whether these macrophage populations contribute to the increased seizure severity observed in FIRE KO mice, we treated FIRE KO mice with PLX3397 (or control) for 7 days to further deplete macrophages. Seizure severity scores were compared between FIRE KO mice and FIRE KO mice treated with PLX3397 (“FIRE+PLX”). Notably, “FIRE+PLX” mice exhibited seizure severity scores like FIRE KO mice, indicating that additional macrophage populations beyond microglia do not contribute significantly to seizure severity in this KA model (**Fig. S1a-c)**. Therefore, consistent with the conclusion from our prior findings^14^, studies in FIRE KO mice reveal that microglia regulate seizure susceptibility in a neuroprotective manner.

### “FIRE” KO mice show exacerbated electrical seizures

Our findings contrast with a recent study that examined seizure susceptibility in a PTZ model using FIRE mice and reported no significant roles for microglia in that context during development^26^. Therefore, to determine whether microglial contributions to seizure susceptibility is limited to chemoconvulsive seizure models, we expanded our studies to an electrical kindling model. Specifically, kindling induces gradual neuronal hyperexcitability through repeated electrical stimulation^45^. If microglia are truly dispensable in seizure regulation, as previously suggested by the recent study^26^, we would expect no differences in kindling progression between FIRE KO and FIRE HET mice. However, when FIRE KO mice were kindled, they required fewer stimulations to get to stage 5 seizures (bilateral clonus with loss of balance) compared to FIRE HET littermate mice (**Fig. 2a-b**). Kindling of FIRE knockout mice also increased the fraction of mice that developed spontaneous recurrent seizures (SRS), leading to more frequent and severe seizures per mouse compared to the FIRE HET mice (**Fig. 2c-d).** Furthermore, while ∼ 30% (2 of 7) of HET mice showed spontaneous recurrent seizures (SRS) after kindling, 100% (4 of 4) of KO mice showed SRS. Since most strains of mice do not develop SRS after kindling^45^, this highlights an important role for endogenous microglia in limiting spontaneous seizure generation. Therefore, our studies demonstrate that FIRE KO mice have exacerbated seizures during both chemoconvulsive and electrical kindling.

**Fig. 2:**
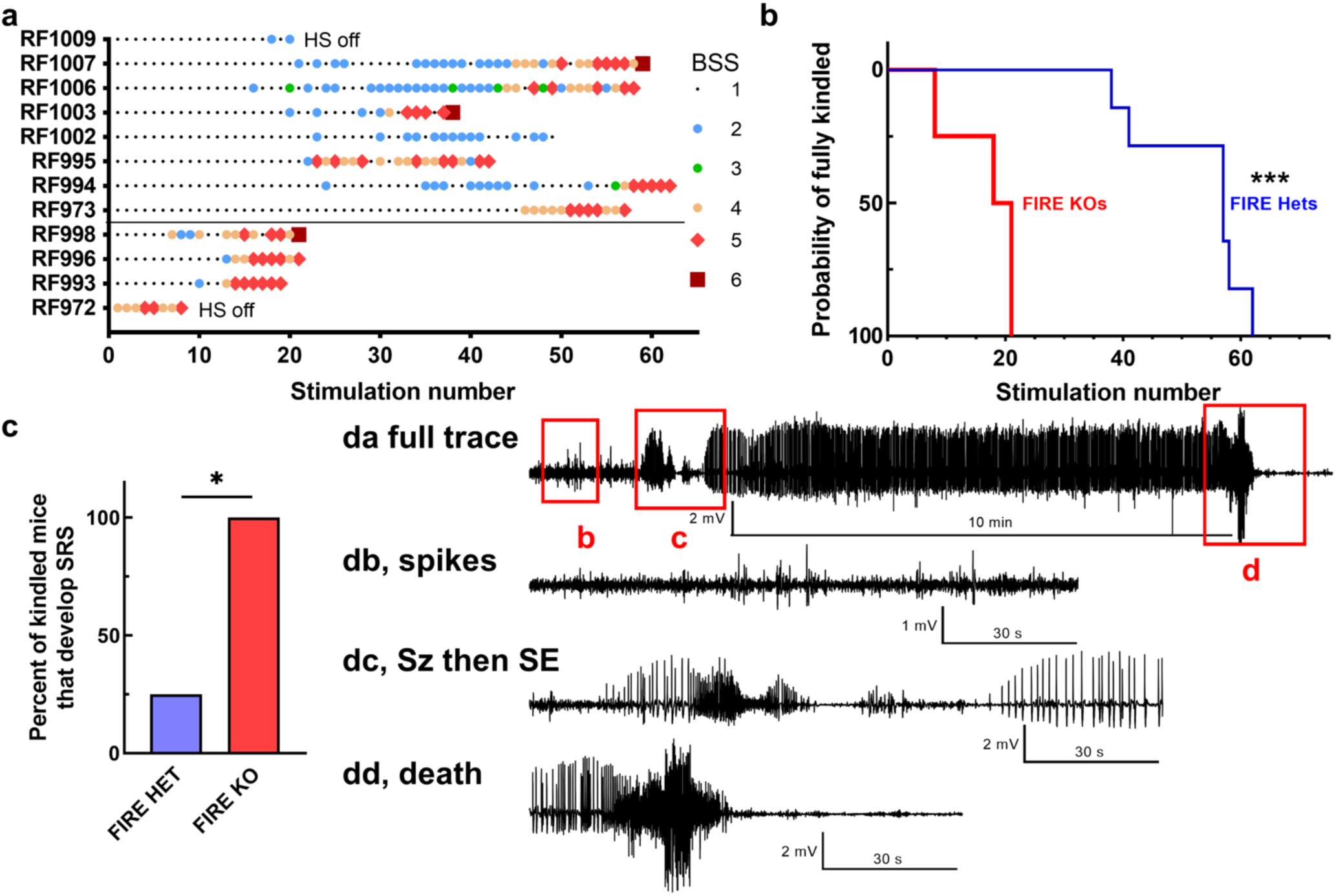
FIRE KO mice are more susceptible to seizures induced by electrical kindling. **a**, Raster plots showing the evolution of the behavior evoked by electrical stimulation of the hippocampus in FIRE HET and FIRE KO littermates. Behavioral seizure score key is shown on the right of panel **a**. “HS off” denotes when the EEG recording headset fell off. Mice are considered fully kindled after electrical stimulation elicits five motor seizures with a BSS score of 5. **b**, Kaplan-Meier plot of the number of electrical stimulations required to reach the fully kindled state in FIRE HET and FIRE KO littermates. **c**, The number of mice that developed spontaneous recurrent seizures after kindling. **d**, Representative EEG traces from a FIRE KO mouse. **da** Shows the final 960 s of EEG from mouse RF974. Red boxes show the regions expanded in **db-dd**. **db** Shows high amplitude spikes in the EEG riding a baseline of high frequency/low amplitude spikes. **dc** Shows an example of the EEG transitioning into a high frequency/high amplitude seizure, followed by a short post-ictal depression. This transitions into a 10-minute-long seizure, which fulfills the definition of status epilepticus (SE). **dd**, SE ends with a discrete seizure and death. N = 4 Hets and 7 KO littermates. Statistical analysis in panel **b** by Mantel-Cox indicates P=0.0005, and panel c by Chi-square, P=0.0143.

### A global or microglial-specific P2RY12 deficiency exacerbates chemoconvulsive kainic acid-induced seizures

Having established that microglia are required for limiting seizure severity in a chemoconvulsive (**Fig. 1**) and an electrical (**Fig. 2**) seizure model using FIRE mice, we next sought to investigate possible mechanisms involved in microglial protection against seizure severity. The purinergic receptor P2RY12 is a microglial-enriched molecule involved in microglial process extensions towards areas of hyperexcitability and can be considered to play an important role in microglial control of seizure intensity^32,42^. In previous work, we had tested the role for P2RY12 in chemoconvulsive seizures using global KO mice^32^. However, in that study, littermates were not used raising possibilities of differences that may result from genetic strains. To overcome this and more conclusively test for a role for P2RY12, we assessed the seizure susceptibility in global P2RY12 knockout (gKO) and wildtype littermates after KA treatment. Seizures persisted in gKO mice compared to their WT littermates over 3h and there was a significantly higher peak seizure score in P2RY12 KO mice compared to WT littermates (**Fig. 3a-b**). Furthermore, the survival curves also showed that there was reduced mortality in gKO mice compared to WT littermates with KA treatment (**Fig. 3c**). These results are consistent with our previous work suggesting roles for P2RY12 in seizure susceptibility^32^.

**Fig. 3:**
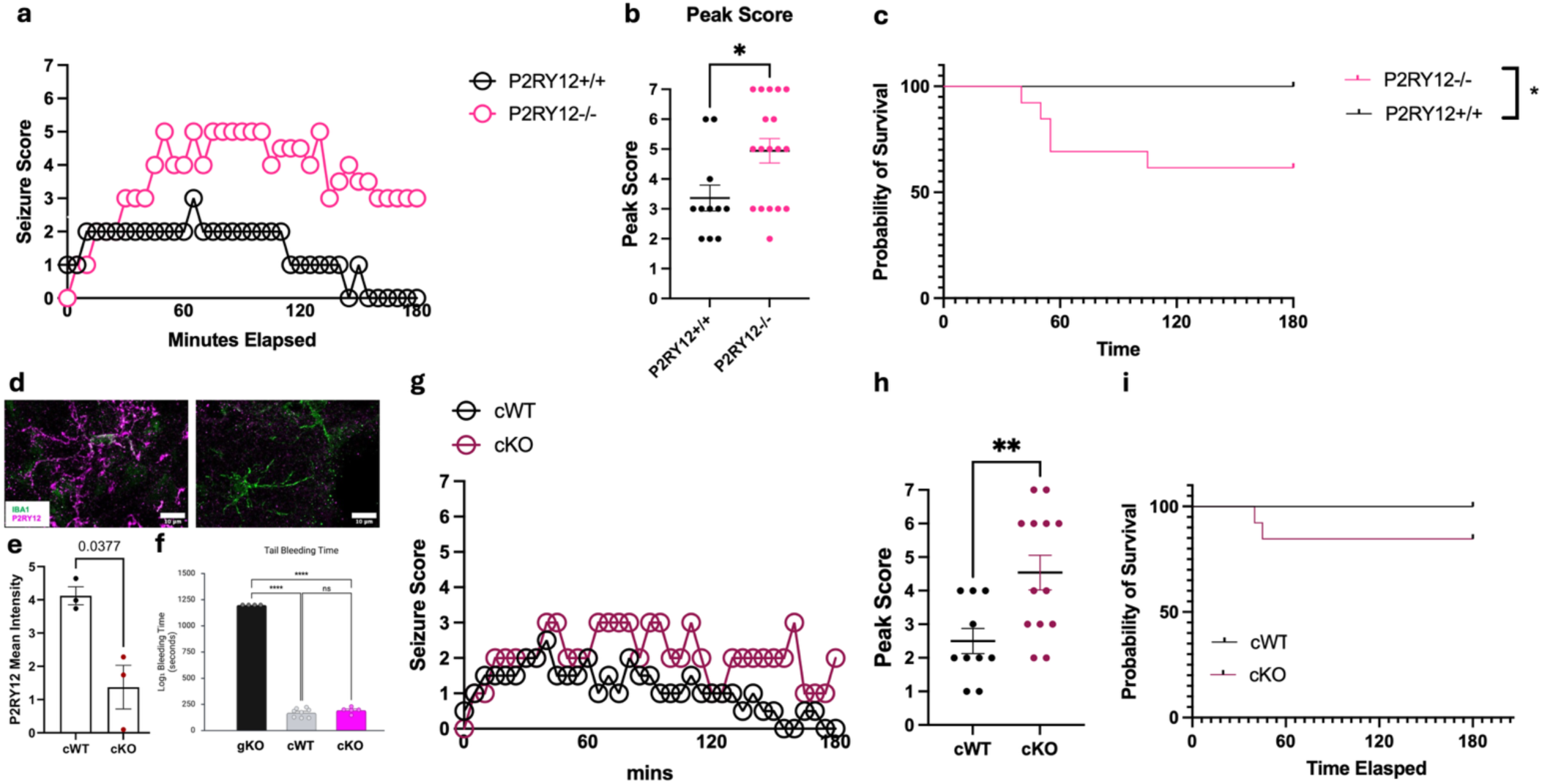
P2RY12 signaling in microglia is required for seizure suppression. **a,** Time course of modified Racine seizure scores in P2RY12-deficient mice. **b**, Quantification of peak seizure score in P2RY12^⁻/⁻^ mice (p < 0.05, unpaired t-test). c, Kaplan–Meier survival analysis of survival probability in P2RY12^⁻/⁻^ mice (p < 0.05, log-rank test). N = 11-18 mice each. **d**, Representative confocal images of IBA1 (green) and P2RY12 (magenta) immunostaining in cWT(P2RY12^fl/fl^:CX3CR1^Cre/+^) and conditional knockout cKO (P2RY12^fl/fl^:CX3CR1^Cre/+)^ mice confirm selective microglial deletion of P2RY12. **e**, Quantification of P2RY12 mean fluorescence intensity in cWT and cKO mice (unpaired t-test). N = 3 mice each. **f**, Tail bleeding assay comparing platelet function across genotypes confirming microglia-specific targeting (p < 0.0001, one-way ANOVA with Tukey’s post hoc test). N = 4-8 mice each. **g-i**, Modified Racine seizure scores overtime of cWT and cKO (**g**), peak seizure score (**h**, p < 0.01), and Kaplan–Meier survival analysis (**i**). N = 10-13 mice each.

Nevertheless, because P2RY12 is also expressed in circulating blood platelets, where it regulates ADP-dependent aggregation and thrombosis^46^, it remained unclear whether the heightened seizure severity observed in global knockout mice resulted from the loss of microglial P2RY12 specifically or from broader systemic effects including for example platelet dysfunction. To resolve this ambiguity, we next utilized a microglia-specific constitutive P2RY12 KO model to determine whether loss of P2RY12 within microglia alone is sufficient to exacerbate seizure outcomes using a microglia-specific conditional knockout mouse model (P2RY12^fl/fl^;Cx3cr1^Cre/+^), hereafter referred to as P2RY12 cKO mice and littermate P2RY12^flf+^;Cx3cr1^Cre/+^ mice, hereafter referred to as P2RY12 cWT mice. In the cKO mice, P2RY12 expression was reduced in microglia (**Fig. 3d-e**). Consistent with a selective loss of P2RY12 loss in microglia without effect on platelets, a bleeding assay on cWT, cKO and gKO mice revealed significantly increased bleeding times only in the gKOs (**Fig. 3f**) indicating preserved platelet P2RY12 function in cKO mice.

Turning to KA-induced seizures, we again report a significantly increased peak seizure scores in P2RY12 cKO mice compared to littermate cWT mice (**Fig. 3g-h**). However, while the seizure severity phenotype for cKOs was consistent with that seen in gKOs, the cKOs did not show significantly different mortality rates compared to their cWT controls (**Fig. 3i**). These data collectively validate the essential role of microglial P2RY12 signaling in regulating neuronal hyperexcitability in response to KA-induced seizures.

### P2RY12 signaling maintains microglial ramification in response to KA

Microglial ramification, the degree of process branching and arborization, is considered a major marker of microglial homeostasis and responsiveness to changes in the surrounding tissue. Ramified microglia are constantly surveying the brain parenchyma by extending and retracting their processes to maintain balance^47,48^. We first conducted microglial tissue surveillance measurements and confirmed that global P2RY12 WT, HET and KO mice (**Fig. S2a**) exhibit similar surveillance indices *in vivo* including their process extension (**Fig. S2b**), process retractions (**Fig. S2c**) and process velocity (**Fig. S2d**) rates. This is consistent with previous reports^36,49^.

Next, to evaluate whether P2RY12 signaling influences microglial structural remodeling during seizure activity, we performed Sholl analyses on microglia in hippocampal CA1 (**Fig. 4a-b**) and the somatosensory cortex (ssCTX **Fig. 4c-d**) regions of fixed tissue from WT and gKO littermate mice under basal and post-KA (90 minutes) conditions. We selected 90 minutes because this seemed like the earliest time when differences between microglial- or P2RY12-sufficient versus deficient mice show seizure severity differences (**Fig. 3a, g**). Under basal conditions, WT and gKO microglia were highly ramified with indistinguishable ramification indices in accordance with microglial surveillance function in both the CA1 (**Fig. 4b**) and ssCTX areas (**Fig. 4d**). Following KA-induced seizures, WT microglia were able to maintain their ramified morphology to a greater extent, whereas there was substantial reduction in process branching in gKO microglia in comparison to WT microglia in both the CA1 (**Fig. 4b**) and ssCTX (**Fig. 4d**) areas. These results indicate that P2RY12 signaling is critical to the preservation of microglial structural complexity in the context of seizure activity. The reduced ramification in microglia from gKO mice underscores increased microglial de-ramification which might lead to diminished structural engagement with hyperactive circuits and therefore contribute to the increased neuronal activation and seizure severity observed our experiments.

**Fig. 4:**
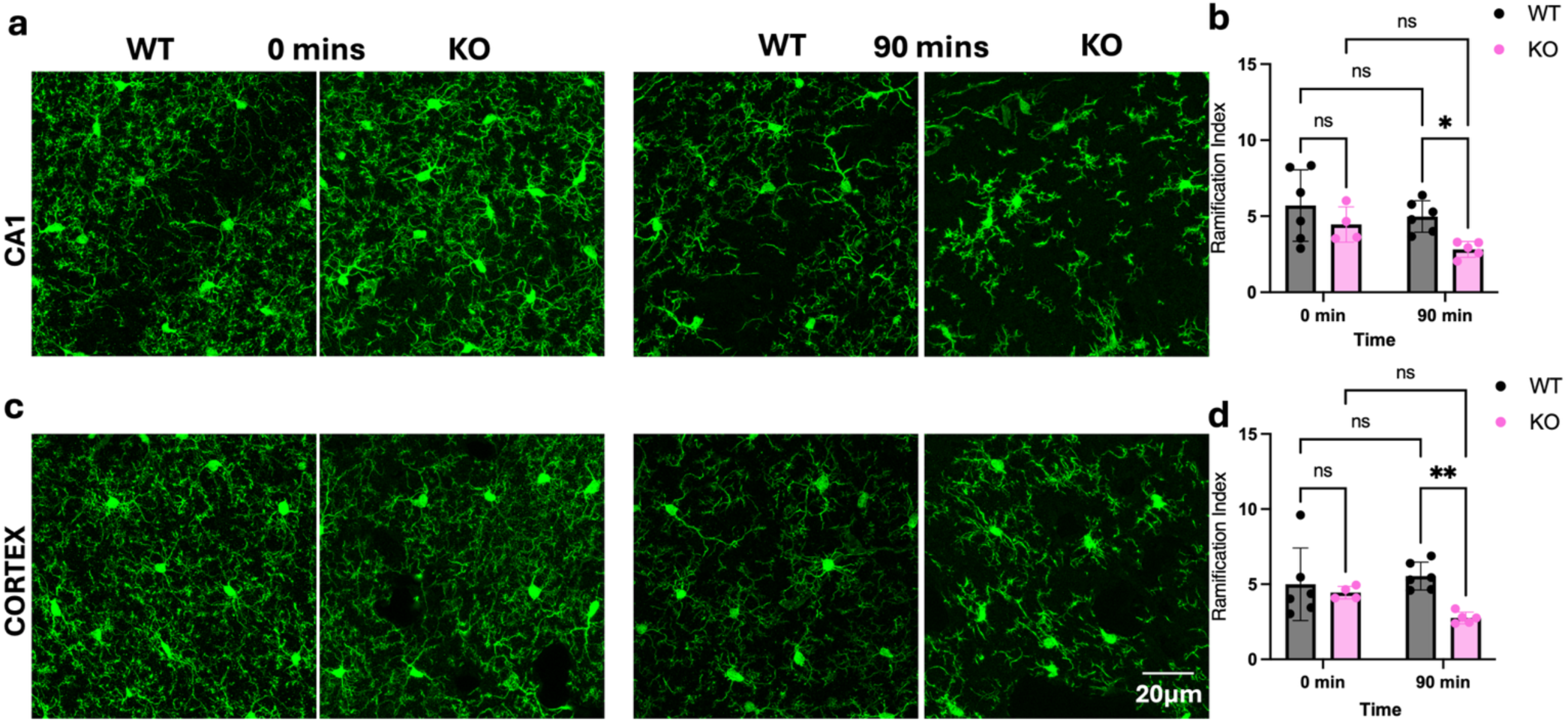
P2RY12 regulates microglial morphological complexity following seizures. **a–d**, Representative confocal images (**a, c**) of IBA1^+^ microglia and quantification of the ramification index (**b, d**) from hippocampal CA1 (**a-b**) and the somatosensory cortex (**c-d**) regions in wildtype (WT) and P2RY12^⁻/⁻^ (KO) littermate mice under baseline (0 min) and post-seizure (90 min) conditions following kainic acid administration. Data shown as mean ± SEM. N = 4-6 mice each. Statistics done by two-way ANOVA with Tukey’s post hoc test.

### A P2RY12 deficiency enhances neuronal activation after KA treatment

As indicated earlier, a P2RY12 deficiency leads to increased seizure severity (**Fig. 3**). Therefore, we further investigated if this increased seizure severity was paralleled by any shifts in neuronal activation corresponding to seizure induction. To address this question, we employed cFos immunolabeling, known to indicate neuronal activity, to evaluate the level of seizure-evoked activation in hippocampal CA1 and the somatosensory cortex of wildtype and gKO mice at basal levels and 90 min of KA treatment. At basal states (0 min), expression of cFos was lower in gKO mice compared to wildtype mice in both brain regions (**Figure 5a-d**). However, 90 minutes post-KA treatment, the P2RY12 gKO mice showed higher expression of cFos than wildtype mice in both brain regions, indicating higher neuronal activity in the absence of P2RY12 signaling during seizures (**Figure 5a-d**). These data indicate that P2RY12 signaling is involved in controlling seizure-induced neuronal excitation, especially in the hippocampus and cortex. The exaggerated cFos response in P2RY12 KO mice suggests that microglial P2RY12 activity is required to limit excessive neuronal firing during KA-induced hyperexcitability, consistent with the increased seizure severity (**Fig. 3**) and impaired microglial ramification (**Fig. 4**) observed in previous studies.

**Fig. 5:**
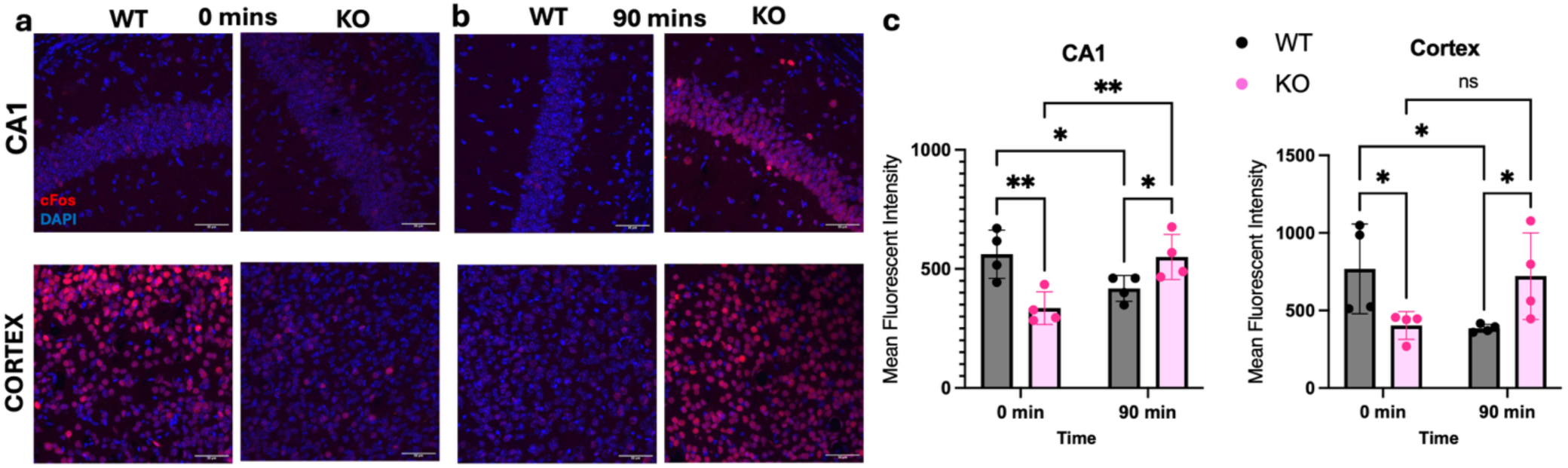
P2RY12 deficiency enhances cFos expression following seizures. **a–b**, Representative confocal images of cFos immunoreactivity from the hippocampal CA1 (**a**) and somatosensory cortex (**b**) regions in wildtype (WT) and P2RY12^⁻/⁻^ (KO) mice under baseline (0 min) and post-seizure (90 min) conditions following kainic acid administration. **c**, Quantification of mean cFos fluorescent intensity in the CA1 (left) and cortex (right) across genotypes and time points. N = 4 mice each. Data shown as mean ± SEM. Statistics by two-way ANOVA with Tukey’s post hoc test.

### P2RY12 signaling maintains inhibitory synaptic coverage during seizures

Since loss of P2RY12 signaling led to exaggerated neuronal activation, we next asked whether this hyperexcitability was accompanied by changes in inhibitory synaptic signaling with a P2RY12 deficiency. To test this, we examined expression of the vesicular GABA transporter (VGAT), a presynaptic marker of inhibitory terminals in both hippocampal CA1 and cortical regions of gKO and wildtype mice at baseline and 90 minutes following KA–induced seizures. Under basal conditions, WT mice showed extensive VGAT area coverage in CA1 and cortex, while gKO mice showed reduced VGAT area even in the absence of seizures, indicating an inherent deficiency in neuronal inhibition under basal conditions. Upon seizure induction, the VGAT area was reduced in both the CA1 and cortex in WT mice, with significant reduction detected only in the cortex. However, in gKO mice in both brain regions VGAT expression was almost non-existent at 90 minutes of KA-treatment (**Fig. 6a-c**). These findings demonstrate that P2RY12 signaling is critical for maintaining inhibitory synaptic coverage in both the absence and presence of seizure activity. The pronounced reduction in VGAT-positive area in gKO mice indicates that disruption of microglial P2RY12 impairs the preservation of inhibitory terminals in basal and hyperexcited states, providing a potential mechanism for the heightened neuronal activation and seizure severity observed in these mice.

**Fig. 6:**
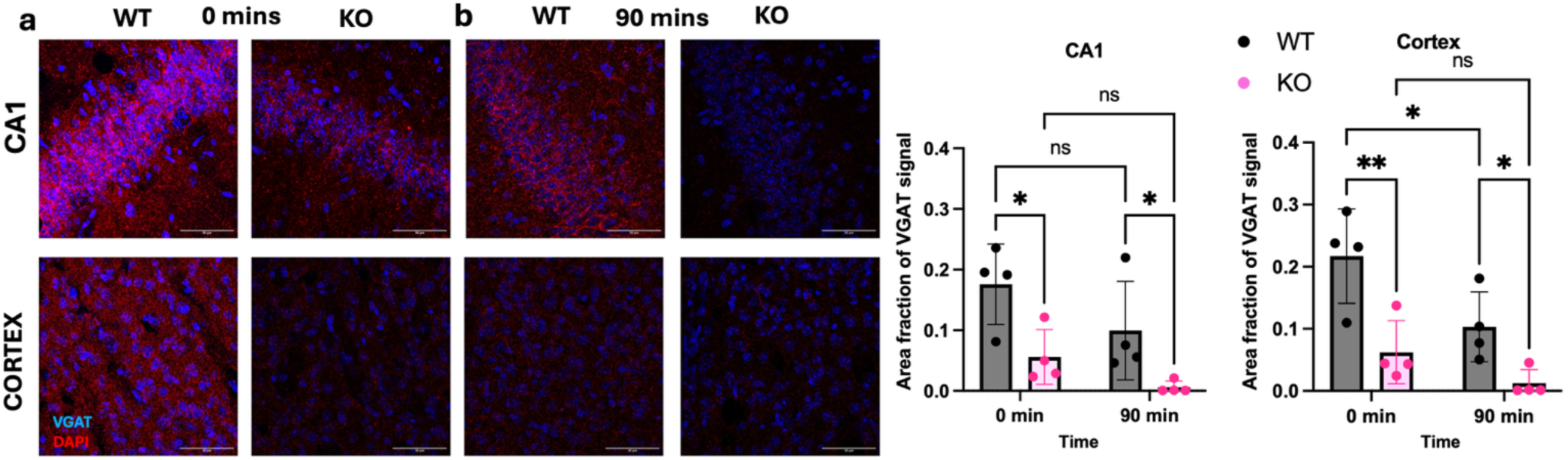
P2RY12 deficiency reduces inhibitory synaptic marker expression following seizures. a–b, Representative confocal images of VGAT immunoreactivity from the hippocampal CA1 (**a**) and somatosensory cortex (**b**) regions in wildtype (WT) and P2RY12^⁻/⁻^ (KO) mice under baseline (0 min) and post-seizure (90 min) conditions following kainic acid administration. **c**, Quantification of VGAT-labeled area per μm of tissue section in the CA1 (left) and cortex (right) across genotypes and time points. N = 4 mice each. Data shown as mean ± SEM. Statistics by two-way ANOVA with Tukey’s post hoc test.

## DISCUSSION

In the current work, we show that mice with microglial-specific genetically deficient mice show increased seizure severity in response to chemoconvulsive kainic acid (KA)-induced seizures (**Fig. 1**) and electrical seizures (**Fig. 2**) consistent with previous results from us and others using pharmacological^15^ and pharmacogenetic^13–15^ approaches. Together, these results provide overwhelmingly supportive evidence for highlighting a neuroprotective role for microglia in various seizure paradigms. Furthermore, both a global and a microglial-specific P2RY12 deletion provided molecular support for microglial contributions to experimental seizures (**Fig. 3**), again consistent with previous report for neuroprotective microglial contributions to seizures from broad Gi^50^ and specific P2RY12^32,42^ mechanisms. Additionally, we find that a P2RY12 deficiency impairs microglial abilities to maintain their morphological state following seizures (**Fig. 4**) co-incident with increased neuronal activation states (**Fig. 5**) and reduced inhibitory synaptic density (**Fig. 6**). In summary, this work shows that microglia are integral to the control of hyperexcitability, that P2RY12 signaling is essential to microglia in controlling excitability, and that P2RY12 has implications for microglial morphology, network activation, and inhibitory synapse structure in networks during seizures.

### Microglial roles in experimental seizures

Microglial roles in seizure disorders have been hotly debated. Initial ideas at the beginning of the century suggested that because of the known seizure-promoting roles for inflammatory mediators like IL-1ß and microglial ability to release such inflammatory molecules, microglia were suggested to perform seizure-promoting functions^51,52^. For example, several studies have shown that following KA-induced seizures, microglia express IL-1ß and other pro-inflammatory cytokines^18–22^. These cytokines facilitate neuronal excitability^53–55^ and thus would promote seizures in such a context. Therefore, by such considerations, it can be expected that microglial activities in seizure contexts promote seizures. Furthermore, in the developing brain, peak seizure sensitivity occurred at the same time as peak microglial density^23^. Nevertheless, while this association was suggested to be causal, evidence provided was at best correlational and lacked direct causal testing.

More recently, several approaches have been employed to more directly test microglial roles in seizure disorders. In an initial pharmacogenetic microglial ablation approach, microglia were found to be inconsequential to acute seizure phenotypes following chemoconvulsive pilocarpine administration. However, the same study showed that following an inflammatory pre-conditioning with LPS, microglia were neuroprotective to pilocarpine induced seizures^13^. Consistent with this, a different pharmacogenetic approach also showed neuroprotective roles for microglia in chemoconvulsive kainic acid-induced seizures^14^. However, such pharmacogenetic approaches have been shown to themselves elicit neuroinflammation^16^ making it difficult to ascertain whether exacerbated seizures with microglial ablation by these methods are a result of losing neuroprotective microglia or having increased neuroinflammation. These concerns raised the need to employ microglial ablation strategies that concomitant inflammation.

In recent years between 2014 and 2019, two such approaches have been developed including a pharmacological approach using the PLX family of drugs to eliminate microglia^40,43^ and a genetic approach with the “FIRE” mice^25^. In a previous study, using the PLX approach, we that microglia perform neuroprotective roles in multiple seizure paradigms^15^. These results suggest that the exacerbated phenotypes from pharmacogenetic approaches likely result from microglial absence rather than other confounds such as inflammation. Like these results, experiments conducted in a Dravet syndrome model (a genetic model of seizures) in zebrafish showed that while microglia increase their inflammatory IL-1ß expression in this seizure model, their ablation using genetic targeting approaches resulted in increased neuronal activity.

Contrary to these findings, a recent study reported, at least in a PTZ chemoconvulsive model during development, that FIRE mice which lack microglia do not show altered seizure severity^26^. However, that study examined only one chemoconvulsive seizure paradigm in which all mice died suggesting that perhaps the induced seizures were too severe to observe differences. Thus, in the current study we employed the FIRE model in adult chemoconvulsive and electrical seizure paradigms and confirmed that a genetic ablation of microglia exacerbates seizures in these paradigms, which is congruent with the majority of studies investigating microglial roles in experimental acute seizure models^13–17^.

### P2RY12 roles in experimental seizures

Molecular mechanisms by which microglia limit seizure severity remains understudied. In the current study, we investigated roles for the P2RY12 receptor in chemoconvulsive KA-induced seizures. Interest in P2RY12 was generated from prior work showing that it is a predominant receptor through which neurons communicate with microglia especially in hyperactive conditions.^28,32,56,57^ Moreover, microglial Gi-signaling dampens neuronal excitability in seizure contexts^50^ and P2RY12 is the most highly expressed Gi-coupled receptor in microglia^27^. Furthermore, previous work in a developmental febrile seizure model^42^ and an adult D1 agonist-induced seizure model,^58^ indicated that microglial P2RY12 mitigate seizure severity. Additionally, we previously showed that a P2RY12 deficiency led to exacerbated KA-induced seizures^32^. However, those studies were not conducted on littermates raising the possibility that genetic differences broadly rather than a P2RY12-specific deficiency could account for the aggravated seizure phenotypes in P2RY12 KO mice. Therefore, to more conclusively explore P2RY12 roles in KA-induced seizures, we examined seizures in littermate mice and confirm more severe seizures in littermates lacking P2RY12. Furthermore, with the development of P2RY12 flox mice^59^, we were able to assess microglial-specific contributions and confirmed the results with global KO mice that microglia’s role is to limit seizure severity.

In addition, an examination of microglial phenotypes revealed an accelerated de-ramification of microglia in P2RY12 KO microglia following KA treatment. Since microglial ramified processes facilitate their tissue surveillance,^47,48^ which regulates microglial physical interactions with neurons such as synapses^60–62^, these results suggest that an increased de-ramification would result in reduced microglial surveillance as well as microglial-neuronal interactions during seizures consistent with previous results.^32,50^ These changes in microglia correlated with increased broad neuronal activity and reduced GABAergic inhibitory tone suggesting that microglial process dynamics under P2RY12 control regulates neuronal hyperactivity during seizures. Downstream mechanisms of P2RY12 regulation of neuronal phenotypes remain to be determined and should be a focus of subsequent work. Nevertheless, our studies herein firmly implicate microglia and its P2RY12 receptors in the regulation of seizure severity during acute seizures warranting future studies.

## MATERIALS AND METHODS

### Animals

All animal experiments were carried out pursuant to the relevant guidelines and regulations of the University of Virginia and approved by the Institutional Animal Care and Use Committee with protocol number 4237-08-21. The animals were housed under controlled temperature, humidity, and light (12:12 hr. light: dark cycle), with food and water readily available *ad libitum*. This study used both male and female mice on a C57BL/6J background between 2-4 months of age, and consisted of the following genotypes: C57Bl/6J mice as wildtype mice; CX3CR1^GFP/+^ expressing GFP under control of the fractalkine receptor (CX3CR1) promoter (Jackson Lab, #005582); P2RY12 KO mice; CX3CR1^Cre^ mice (Jackson Lab, #025524); P2RY12^fl/fl^ mice as a generous gift from Dr. Long-Jun Wu at the University of Texas Health in Houston; *Csf1r^ΔFIRE/ΔFIRE^*as a generous gift from Dr. Sandro da Mesquita, Mayo Clinic. Mice were housed in groups for the experiment without special environmental enrichment.

### Chemoconvulsive seizure induction

For chemoconvulsive seizure initiation model, 2-4-month-old mice were used, and i.p. injected with kainic acid (KA) at 24mg/kg body weight. Control mice were injected with equal volumes of saline that was used as the vehicle to dissolve the kainic acid. Seizures were scored using a modified Racine scale as follows: (1) freezing behavior; (2) rigid posture with raised tail; (3) continuous head bobbing and forepaws shaking; (4) rearing, falling, and jumping; (5) continuous occurrence of level 4; and (6) loss of posture and generalized convulsion activity ^22,34,63^. Scores were recorded every 5 minutes for up to 3 hours. Scores were given based on the summary of the mouse behavior over the 5-minute interval. The experimenter was “blind” to the prior condition of the mice. The seizure scores are presented as the median of the scores and the area under the curve (AUC) was analyzed. For pharmacological depletion, mice were treated with PLX3397 (660mg/kg) for 7 days prior to KA treatment.

### Animal surgeries

Surgeries were performed on 8-week-old mice using isoflurane anesthesia. A Kopf stereotaxic apparatus was used to guide implantation of an EEG recording headset that included a bipolar Teflon-coated stainless-steel stimulating electrode (A-M Systems, diameter = 0.008”, #791400), bilateral supradural cortical electrodes, and a cerebellar reference electrode. Electrodes were connected to a six-pin pedestal (Plastics One), which was secured to the skull with dental cement. The depth electrode was placed in the left hippocampus, targeting the perforant path at the following coordinates (from bregma): 3 mm posterior, 3 mm lateral, and 3 mm depth. Animal discomfort was minimized with bupivacaine and ketoprofen (4 mg/kg delivered subcutaneously after surgery and every 24 hours as needed). All studies were performed in accordance with protocols approved by the Animal Care and Use Committee of the University of Virginia and ARRIVE guidelines.

### Kindling protocol

After at least 1 week for recovery from surgery, mice were connected to a video-EEG monitoring system (AURA LTM64 using TWin software, Grass) via a flexible cable and commutator (Plastics One). One day later, we measured the afterdischarge threshold (ADT); the hippocampal electrode was connected to a constant current stimulator (A-M Systems, Model 2100), and then a 1-millisecond biphasic square wave pulse at 50 Hz was applied for 2 seconds. The current was initially set at 20 μA and increased in 20-μA increments until an electrographic discharge was observed (2 minutes between stimulations). The mean ADT was 108 μA (range = 40-400). For kindling, the current intensity was set to 1.5 × the magnitude of ADT for that mouse and was delivered twice per day, every other day, with an interval between stimulations of at least 4 hours. Animals were considered fully kindled when stimulations evoked five consecutive seizures with a behavioral score of at least 5 (bilateral clonus with loss of posture control) on a modified Racine scale. Animals were monitored by continuous video-EEG throughout the experiment (24 h/d, 7 d/wk). EEG data were digitized at 400 Hz, then stored on a file server along with video for subsequent analysis. Spontaneous seizures were defined as evolving spike-wave discharges with >2-Hz frequency and 3 × the baseline amplitude lasting 15 seconds or longer. During analysis, electrographic seizures were confirmed by the corresponding video and a behavioral score was assigned.

### Immunohistochemistry and confocal microscopy

Mice euthanized perfused with 1x PBS (Life Technologies Corporation, NY, USA) + Heparin 0.05% prior to brain harvesting and post-fixation in 4% PFA for 24 hours, with the exception of brains used for c-Fos staining, which were only post-fixed in 4% PFA for 1-2 hours. Brains were then washed in PBS and placed in 15% sucrose solution at 4°C for 24 hours, then transferred to 30% sucrose solution at 4°C for 24 hours. Brains were then cryopreserved in optimal cutting temperature compound (OCT) media (Product #: 72592; Electron Microscopy Science, USA) and then sectioned into 40-µm-thick free-floating sections obtained by sectioning brains with a cryostat (Model CM1950, Leica). Four sections per mouse were incubated in blocking buffer consisting of 1% Bovine Serum Albumin (BSA, Sigma-Aldrich, Germany, 5% normal donkey serum (Product # 017-000-121; Jackson ImmunoResearch Laboratories Inc. USA) and 0.3% Triton X-100 solution (Product #: 93443; Sigma-Aldrich) in 1x PBS for 1 hour at room temperature. Sections were then immersed in primary antibody and blocking buffer solution and placed on a shaker overnight at 4°C.

Primary antibodies used were as follows: rabbit anti-Iba1 (1:500; Cat #019-19741; FUJIFILM Wako Pure Chemical Corporation), rabbit anti-cFos (1:1000; Cat #226008; Synaptic Systems), rabbit anti-VGAT (1:1000; Cat #131004; Synaptic Systems). All secondary antibodies were used at 1:500 dilution and were incubated in 1x PBS, 5% normal donkey serum, and secondary antibodies for 2 hours in the dark. Secondary antibodies used include the following: donkey anti-guinea pig IgG Alexa Fluor 647 (Cat #706-605-148; Jackson ImmunoResearch Laboratories Inc.), donkey anti-rabbit IgG Alexa Fluor 488 (Cat #A-21206; ThermoFisher Scientific), goat anti-rabbit IgG Alexa Fluor 488 (Cat #A-11034; ThermoFisher Scientific), goat anti-rat 647 IgG Alexa Fluor 647 (Cat #A-21247; ThermoFisher Scientific), goat anti-rat 680 IgG Alexa-Fluor 680 (Cat #A21096; ThermoFisher Scientific), donkey anti-rat IgG Alexa Fluor 647 (Cat #712-605-153; Jackson ImmunoResearch Laboratories Inc.), donkey anti-rabbit IgG Alexa Fluor 594 (Cat #R37119; ThermoFisher Scientific), donkey anti-rabbit IgG Alexa Fluor 647 (Cat #A-31573; ThermoFisher Scientific), goat anti-mouse 647 IgG Alexa Fluor 647 (Cat #A32728; ThermoFisher Scientific), and goat anti-mouse IgG Alexa Fluor 488 (Cat #A-11001, ThermoFisher Scientific). Cell nuclei were stained by incubating sections in NucBlue (1 drop per 1 mL of 1x PBS; Hoechst 33342; ThermoFisher Scientific). Slides were imaged on a Stellaris 5 confocal microscope (Leica Microsystems, USA) with 20x, 40x, and 63x objectives.

### Sholl analysis of microglial morphology for ramification index

The ramification of microglial processes was determined using a Sholl analysis through the SNT plugin in ImageJ. Slices including the hippocampus (the CA1 region) and the cortex were stained for Iba1 to identify microglia. Microglia from 3-5 non-overlapping fields of view and displayed well-defined bodies and processes were randomly selected from each mouse. After using a custom segmentation plugin with Ilastik, the images were converted to 8-bit grayscale and a binary threshold to identify individual cells. The Sholl analysis plugin plotted concentric circles 1μm apart and determined the number of microglial branch intersection points for each circle. The Sholl analysis parameters of total intersection and the ramification index were determined from exported datasets analyzed in R. The data was analyzed using a one-way ANOVA after checking residuals to determine difference between groups.

### In vivo multiphoton imaging

To perform cranial window surgery, induction of surgical plane anesthesia with 2-5% isoflurane was first established. Pre-operative analgesics (Bupivacaine) were then administered subcutaneously at the site of incision prior to surgery. Hair and skin of the skull were then removed, and a 3 mm craniectomy 2 mm posterior and 1.5 mm lateral to bregma performed on one hemisphere on the somatosensory cortex. Dura was then removed, followed by the placement of a 3 mm circular No. 1 cover glass (Warner Instruments CS-3R, Cat #: 64-0720). A light-curing dental cement (Tetric EvoFlow) was applied and cured with a Kerr Demi Ultra LED Curing Light (DentalHealth Products). iBond Total Etch glue (Heraeus) was applied to the rest of the skull, except for the region with the window. This was also cured with the LED Curing Light. The light-curing dental glue was used to attach a custom-made head bar onto the other side of the skull from which the craniotomy was performed. Mice were allowed to recover for at least 2 weeks before imaging. To image, CX3CR1^GFP/+^ mice were placed on a Kopf sterotax with heating pad and were anesthetized with isofluorane (1.5%). Imaging was performed 0-200 µm below the surface of the somatosensory cortex. Optical sections were acquired using a Leica SP8 multiphoton microscope with a coherent laser. A wavelength of 880 nm was used to image microglia, and images were collected at a 1024 x 1024 pixel resolution using a 25 x 0.9 NA objective with a 1.5x optical zoom. Several fields of view of z-stack images were collected through a volume of tissue. To observe microglial dynamics, z-stack time-lapse images were acquired every minute at 2 µm steps. Z projections of longitudinal multiphoton videos were created and analyzed using Fiji.

### Statistical analysis

Data were initially measured for normality and homoscedasticity and upon comparing normal distributions and variances further analyzed with the respective tests. Student’s t-test was used to compare two-groups and one-way ANOVA used to compare more than two groups. Other specific tests are stated in the figure legend for each of the experiments.

## Acknowledgments

We thank members of the Eyo Lab, the Center for Brain Immunology and Glia (BIG) and the UVA Neuroscience and Pharmacology Departments for valuable discussions and insights in the development of this project. This study received the following support: National Institute of Neurological Disorders and Stroke, NIH NS119243 (U.B.E) NS112549 (E.P.R), 5T32GM148379 (S.G-S), Gilliam award from the HHMI (S.G-S), and Owen’s Family Foundation (U.B.E).

## FIGURES AND FIGURE LEGENDS

**Fig. S1:**
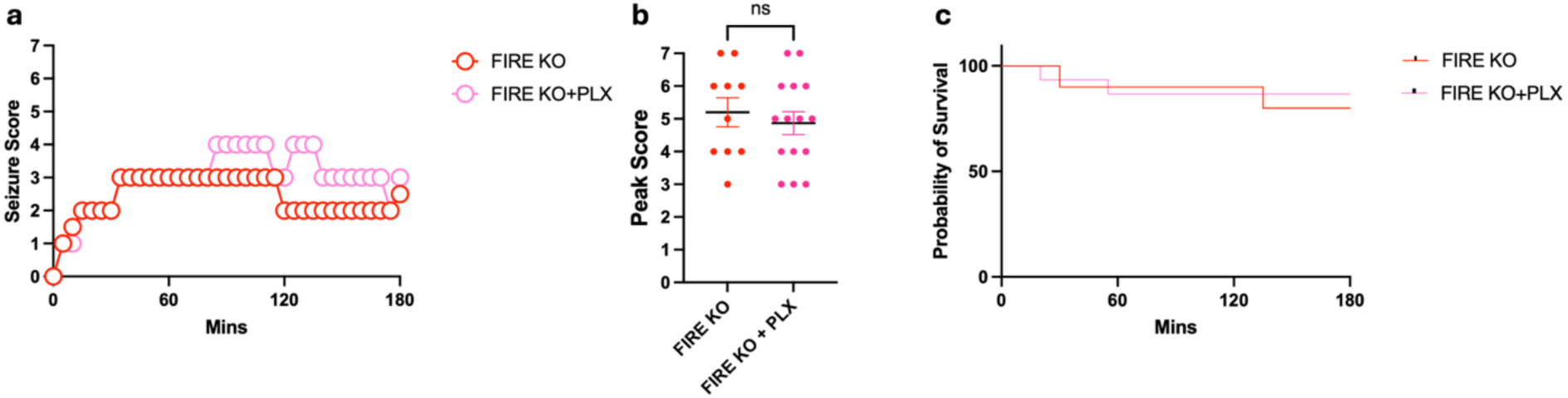
Elimination of border associated macrophages in FIRE KO mice does not alter seizure severity. **a-c**, Seizure severity scores over time in mins (**a**), peak seizure scores (**b**) and (**c**) Kaplan-Meier survival curves of FIRE KO mice and FIRE KO mice with PLX3397 (PLX, 660mg/kg) for 7 days prior to kainic acid treatment. N = 10 - 15 mice each. Data shown as mean ± SEM. Statistics assessed by an ANOVA.

**Fig. S2:**
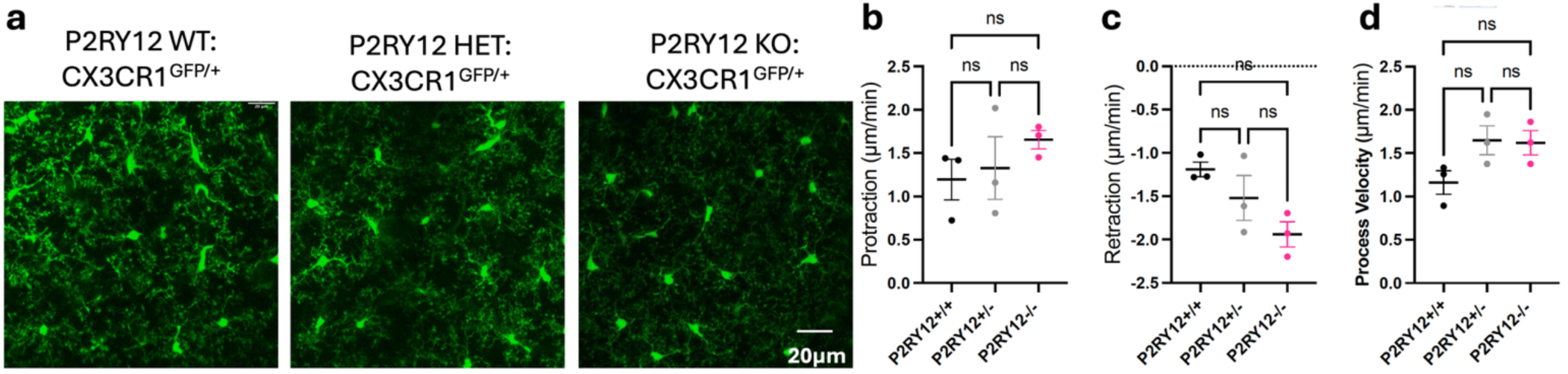
P2RY12 does not regulate microglial tissue surveillance. **a-d**, Representative *in vivo* two photon images of microglia from P2RY12 WT, HET and KO littermates (a) used to quantify process protraction (b), retraction (c) and velocity (d). N = 3 mice each. Data shown as mean ± SEM. Statistics assessed by an ANOVA.

