## Supplementary figures and images for "Microglia and its P2RY12 Receptors Regulate Seizure Severity"

### Supplemental Figure 1

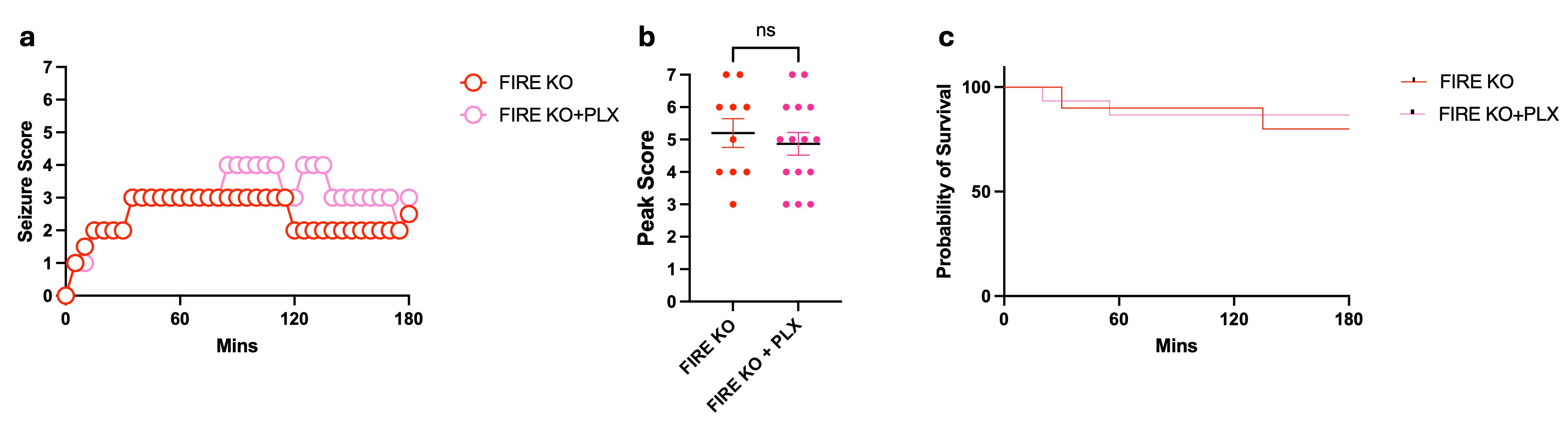

### Supplemental Figure 2

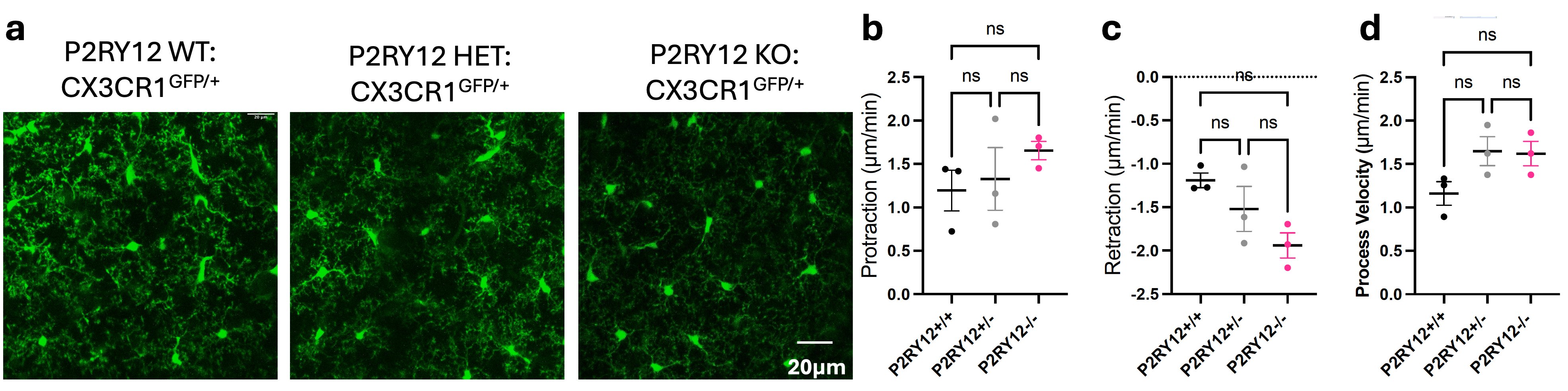
